# Boosting wheat functional genomics via indexed EMS mutant library of KN9204

**DOI:** 10.1101/2022.12.05.519108

**Authors:** Dongzhi Wang, Yongpeng Li, Haojie Wang, Yongxin Xu, Yiman Yang, Yuxin Zhou, Zhongxu Chen, Yuqing Zhou, Lixuan Gui, Yi Guo, Chunjiang Zhou, Wenqiang Tang, Shuzhi Zheng, Lei Wang, Xiulin Guo, Yingjun Zhang, Fa Cui, Xuelei Lin, Yuling Jiao, Yuehui He, Junming Li, Fei He, Xigang Liu, Jun Xiao

## Abstract

A better understanding of wheat functional genomics could facilitate the targeted breeding for agronomic traits improvement and environmental adaptation. With the release of reference genomes and extensive re-sequencing data of wheat and relatives, wheat functional genomics enters a new era. However, limited transformation efficiency in wheat hampers in-depth gene functional study and genetic manipulation for breeding. Here, we generated an EMS mutagenesis library of KN9204, a widely grown elite wheat variety in northern China, with available reference genome, transcriptome, and epigenome of various tissues. The library harbors enormous developmental diversity covering important tissues and transition stages. Exome capture sequencing of 2,090 mutant lines, with probes designed by KN9204 genome, revealed that 98.79% of coding genes have mutations and 1,383 EMS-type SNPs per line averagely. Novel allelic variations for important agronomic trait-related genes, such as *Rht-D1, Q, TaTB1*, and *WFZP*, were identified. We tested 100 lines with severe mutations in 80 NAC TFs under drought and salinity stresses, and found 13 lines with altered sensitivity. Three lines were further analyzed for the regulation insight of NAC TFs in stress response by combing transcriptome and available chromatin accessibility data. Hundreds of direct targets of NAC with altered transcriptional pattern in mutant lines under salt or drought stress induction were identified, including *SNAC1, DREB2B, CML16* and *ZFP182*, knowing factors in abiotic stresses response. Thus, we have generated and indexed KN9204 EMS mutant library which would facilitate functional genomics study and provide resources for genetic manipulation in wheat.

## Introduction

Common wheat is a vital staple crop since the dawn of civilization. Wheat production needs to be significantly increased to feed the ever growing population (Hickey et al., 2019). Understanding the genetic basis and molecular regulation of wheat productivity, end-use quality and environmental adaptability have lagged far behind other cereal crops (Xiao et al., 2022). One of bottleneck is the lack of genetic resources meeting robust and efficient characterization and validation for functional genomics study. Besides, the breeding process have narrowed the available genetic variability of wheat germplasm resources (Liang et al., 2021; Pour-Aboughadareh et al., 2021). Over the past decades, induced mutagenesis has played a vital role in gene discovery, functional characterization and breeding in plants. For example, the genome-wide T-DNA insertion mutagenesis pools has significantly promoted studies of functional genomics in *Arabidopsis* and rice (Krysan et al., 1999; Jeong et al., 2002). Artificial mutagenesis has accelerated breeding, producing trait-improved wheat varieties worldwide, including Ug99-resistant wheat variety ‘Eldo Ngano 1’ and salt tolerant cultivar H6756 from γ-ray mutation (Forster, 2014), high-yielding, sprouting- and lodging-resistant wheat Luyuan502 from universe ray-induced mutation (Liu et al., 2021). Thus, introducing novel sequence variation into crop genomes by artificial mutagenesis is an option to widen the genetic diversity for functional analysis and crop breeding.

Among the physical and chemical mutagenic sources, ethyl methylsulfonate (EMS) has been widely adopted in plants. EMS produces random point mutations, mainly in the form of G/C to A/T transversions (Henry et al., 2014). Compared to physical rays, Ac/Ds and T-DNA insertion mutagenesis, EMS produce a range of mutation strengths, with strong mutations necessary for gene function studies and weak mutations suitable for breeding improvement. Owing to the hexaploid nature, wheat allows for accumulating a higher mutational load without disturbing its survival. This makes wheat easy to generate saturate mutations covering the whole coding genome. For the past years, genome editing tools in particular CRISPR/Cas9, could introduce mutation at specific location, opens an era for studying gene function and precision crop breeding (Gao, 2021). However, genome editing is limited to a few wheat varieties due to the low efficiency of genetic transformation (Wang et al., 2017a). Besides, extensive commercialization of genetically modified crop was restricted due to the current regulatory hurdles in many countries (Palan et al., 2021). Therefore, as a non-transgenic genetics approach without limitation of varieties, EMS-induced mutants were expected to coexist with CRISPR/Cas9 for crop improvement and contributes to the functional genomics research in wheat.

Several EMS mutagenesis populations have been established in a range of wheat cultivars, playing significant roles in expanding breeding germplasms, generation of novel allelic variations, identification and functional study of important regulators. For instance, herbicide-resistant wheat lines were isolated from screening of mutant pools of “Ningchun 4”, “Aikang 58”, “Lunxuan 987”, “Zhoumai 16” and “Jing RS801” (Chen et al., 2021b). Allelic variations in genes *ADP-glucose pyrophosphorylase (AGP) large subunit* (*AGPL*) (Guo et al., 2017), *Starch branching enzyme IIa* (*SBEIIa*) (Botticella et al., 2011; Rawat et al., 2019), *Waxy* (Rawat et al., 2019), *Sucrose transporter* (*SUT1*) (Rawat et al., 2019) and wheat *GA 20-oxidase* gene (*TaGA20ox1*) (King et al., 2015) were screened and used for functional study and breeding. Yet, very few genes have been cloned and characterized from EMS mutagenesis populations. One of the primary reasons is the lack of gene-indexed mutations at a genome-wide scale. Due to the huge size of the wheat genome, it is economically infeasible to obtain variation in large numbers of individuals by whole-genome sequencing (Uauy et al., 2017; IWGSC et al., 2018). Targeted capture and sequencing of coding regions, with less than 2% high-value genomic regions enriched for functional variants and low level of repetitive region, was a viable strategy to identify mutations in wheat (Uauy et al., 2017). Indeed, <u>W</u>hole-<u>E</u>xome capture followed by <u>S</u>equencing (WES) has been widely used in crops with large genome, due to its robust, rapid, cost effective, and high-throughput nature (Kaur and Gaikwad, 2017). Krasileva and colleagues sequenced the coding regions of 1,535 Kronos (tetraploid) and 1,200 Cadenza (hexaploid, spring wheat) mutant lines using an 84-Mb whole-exome capture assay and established *in silico* functional genomic resources in wheat (Krasileva et al., 2017). Nonetheless, Cadenza mutations did not cover all the annotated genes, and more sequence-indexed mutants are still needed. Besides, more mutagenesis library with diverse genetic background are highly demanded for the functional genomic study and breeding, in particular of winter wheat which contribute 70% of the total wheat acreage (Kadar et al., 2018).

In this study, we generated an EMS-mutagenesis library in elite winter wheat variety KN9204 and identified 2.89 million EMS-type SNPs covering 98.79% of high confidence genes through WES. The abundant morphological diversity and novel allelic on agronomic-related genes provide a valuable genetic resource for gene identification and breeding application. Importantly, integration KN9204 mutant lines with our previously generated multi-omic data in KN9204, we take NAC TFs as example to show the robustness for functional genomics study.

## Result

### Generation of EMS-mutagenesis library in wheat cultivar KN9204

Kenong 9204 (KN9204) is a representative winter wheat cultivar widely grown in north China plain, characterized by semi-dwarf (∼ 74 cm), compact plant architecture (∼27,5000 fertile spikes ha^-1^), high yield (∼ 7,500 Kg ha^-1^), and high nitrogen use efficiency (NU_P_E≈65%, NU_T_E≈35%) (Shi et al., 2022; Cui et al., 2016; Jia et al., 2006). Dozens of cultivars derived from KN9204 with improved grain yield, overwintering ability, heat resistance and milling quality have been authorized (**Figure 1A**) (Zhao et al., 2015; Li et al., 2021b). Among them, with moderate regeneration efficiency, Kenong199 is generally used as a typical winter variety in genetic transformation for gene functional study (Wang et al., 2017a). Besides, numerous QTLs of various traits including flag leaf size, spike-related traits, root morphology, grain number and nitrogen use efficiency have been mapped via KN9204-developed bi-parental genetic populations by us and others previously (Fan et al., 2018; Fan et al., 2019; Liu et al., 2020b; Yu et al., 2022; Zhao et al., 2022). Importantly, reference genome (Shi et al., 2022), transcriptome and epigenome including chromatin accessibility, core histone modifications and variants from multiple tissues, developmental stages and environmental treatments have been generated for KN9204 (**Figure 1B, Figure S1**) (Li et al., 2018; Liu et al., 2020b; Liu et al., 2020b; Shi et al., 2022; Lin et al., 2022). Thus, the elite agronomic traits, the clear and productive genealogies, the multi-dimensional omic information including genome, transcriptome, epigenome and the preliminary mapped QTL loci make KN9204 an ideal choice for functional genomic study and breeding improvement via mutagenesis.

**Figure 1.**
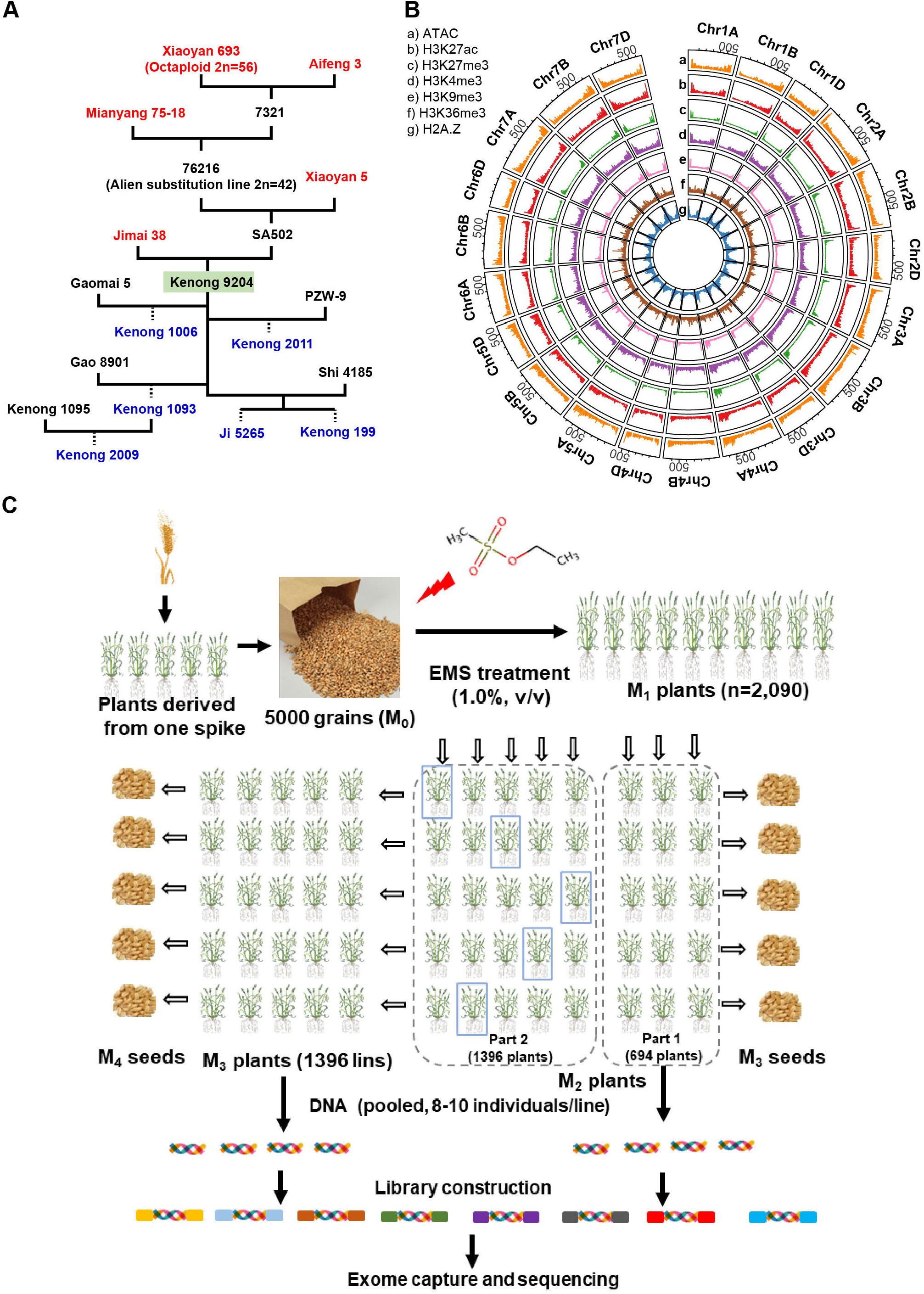
Construction of EMS mutagenesis in winter wheat variety KN9204. (A) A simplified pedigree showing the original parents (in red) and offsprings (in blue) of Kenong 9204 (KN9204). The indirect parents and other cultivars were in black. (B) Circos plot summarizing the chromosomal distribution of multi-epigenomic data. From the outer to the inner circle is: a. ATAC, b. H3K27ac, c. H3K27me3, d. H3K4me3, e. H3K9me3, f. H3K36me3, g. H2A.Z. Bar plots in each circle present the density of epigenetic marks intensity. (C) Overview of the construction and whole exome capture sequencing procedure of EMS mutagenesis population of KN9204.

Accordingly, we generated a mutant library in KN9204 background by treatment of 1.0% v/v EMS. Approximately 5,000 seeds (M_0_) were used for mutagenesis, 2,134 germinated and resulting in M_1_ plants. In total, 2,090 M_1_ plants survived and generated the M_2_ individuals. Among these, 1,396 M_2_ plants successively advanced to the M_3_ generation through single-seed descent with selfing (**Figure 1C**). Sixteen progenies of the 694 M_2_ and 1,396 M_3_ individuals were planted in a row in the field for phenotypic investigation, eight to ten individuals were pooled for DNA extraction and subsequently exome-capture sequencing (**Figure 1C**). Seeds of each mutant line were harvested and bulked for later distribution and further functional study (**Figure 1C**).

Therefore, we have generated a mutant library in elite cultivar KN9204 background by EMS mutagenesis for facilitating functional genomic study and expanding the genetic allelic resources for breeding.

### Diverse developmental variations in KN9204 mutant library

Next, we evaluated the genetic diversity generated in KN9204 mutant library for developmental traits throughout the life cycle of wheat. Two independent surveys were conducted during the 2021– 2022 cropping seasons in the fields at Beijing (39°55’ N, 116°23’ E) and Shijiazhuang (38°04’ N, 114°28’ E) in China. Anomalous developmental defects were observed in both fields and cataloged, including chlorisis, procumbent, twisted leaves or tufted plant at seedling stage (**Figure 2A**); plant architecture alteration such as plant height (PH), tiller number, tiller angle and flowering time variation during heading stage (**Figure 2B**); spike-related traits such as spike length (SL), spikelet density, paired spikelets, degenerated spikelets, florets number, and altered glume hair and awns (**Figure 2C**); leaf morphology (**Figure 2D**) and fertility alteration during grain filling stage, and grain-related traits such as grain number, grain size, grain plumpness, seed coat color, and endosperm hardness (**Figure 2E**) at the mature stage. In total, 286 lines (13.68%) repetitively in both fields exhibited a noticeable alteration in qualitative indicators, categorized into four classes including plant architecture, spike and affiliated organs, leaf shape and color, and other terms. Plant architecture is the largest category with 123 lines, and most of which are PH related mutants (76.42%, 94 out of 123 lines). Spike-related traits are the second most variable characteristics with 114 lines, of which paired-spikelets is presence dominantly (81.57%, 93 out of 114 lines), followed by altered auxiliary organs such as awn and glumes (**Table S1**). Besides, premature senescence, wax leaf and early or late heading are also commonly observed, with more than ten lines each in the KNN9204 mutant library (**Table S1**).

**Figure 2.**
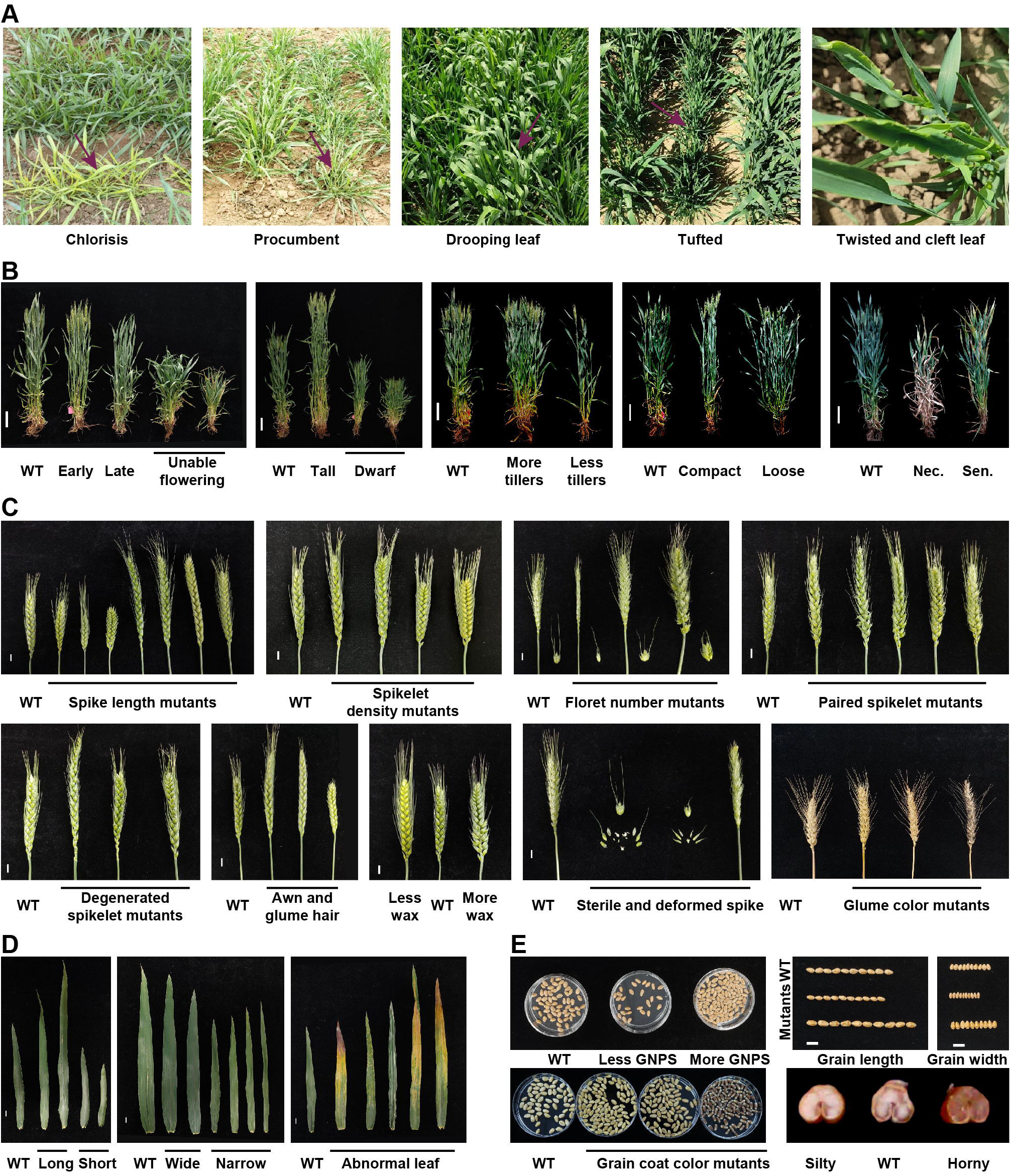
Developmental diversity of various categories within KN9204 mutant library. (A) Anomalous developmental defects observed at seedling stage. The arrow indicates the line with anomalous developmental defects. (B) Plant architecture alteration mutants during heading stage. WT means KN9204 wild-type, Scale bar = 10 cm. Nec., premature necrosis; Sen., premature senescence. (C) Pictures of spike and affiliated organs mutants during grain filling stage. Scale bar = 1 cm. (D) Leaf shape mutants and abnormal leaves, including lesion mimic, striped leaf and premature senescence, etc. Scale Bar = 1 cm. (E) Grain-related mutant phenotypes. Grains from one single main spike of mutants were compared with that of WT KN9204. GNPS, grain number per spike. Scale bar = 1 cm.

To estimate the phenotypic distribution of important agronomic-related traits in KN9204 mutant library, we investigated thirteen quantitative indicators for a randomly chosen 200 mutant lines, including PH, SL, spike number per plant (SNPP), spikelet number per spike (SNPS), grain number per spike (GNPS), fertile florets per spikelet (FFPS), thousand-grain weight (TGW), etc (**Table S2**). Among the quantitative indicators, SL, SNPS, GNPS, FFPS and TGW fit an approximate normal distribution (**Figure S2A-F**). For instance, the mutant population produced slightly reduced SNPS as compared to KN9204 averagely (Mutants versus KN9204, 23.67 versus 25.54) (**Figure S2B**), with a minimum and maximum SNPS of 17.25 and 34.62, respectively (**Table S2**). However, PH tends to fit a positively skewed distribution, as 46 lines with PH more than 80 cm and only six lines less than 60 cm (**Figure S2G**). Grain width and grain roundness is just the opposite, fitting a negatively skewed distribution (**Figure S2H, S2I**).

The high frequency of visible morphological alterations and the wide range of phenotypic variations observed in KN9204 mutant library implied that it would be a valuable resource for screening mutant lines that would aid gene functional study and breeding application.

### Characterizing the EMS-mutagenesis in KN9204 mutant library by WES

To fill the gap between genome and “gene-ome” in hexaploid wheat, we conducted the WES for the EMS-mutagenesis library together with KN9204 wild type (WT), with probes that designed according to the reference genome of KN9204 we assembled recently (Shi et al., 2022) (See methods for details). Averagely 91.62 million high-quality clean reads (Q30 > 0.85) per sample were generated in WES, covering 99.93% of exome targets with an average depth of 11.63-fold as aligned to the KN9204 reference genome (**Table 1**). Based on the alignment, approximately 57.59% of the target region was sequenced at least five times, whereas 41.57% had a more than 10-fold sequencing depth (**Figure S3A**). A minimum coverage of ten mutated reads for heterozygous (HetMC10) and six for homozygous (HomMC6) mutations were chosen to differentiate real mutations from sequencing errors (**Figure S3B**). Using a modified <u>G</u>enome <u>A</u>nalysis <u>T</u>ool<u>K</u>it (GATK) bioinformatics pipeline followed by strict quality filtering (see Methods), we identified a total of 2.97 million SNPs, with homozygous/ heterozygous ratio of 1.83 and 2.74 for the M_2_ and M_3_ population, respectively (**Figure 3A**). As expected, 97.26% SNPs are GC > AT transitions (**Figure 3B**), as EMS preferentially alkylates guanine to O_6_-ethylguanine, leading to the mispairing of G > T during DNA replication (Henry et al., 2014). Consequently, a set of 2.89 million EMS-type SNPs were identified, with estimated average of 1,383 EMS-type mutations per mutant line and 19.94 mutations per Kb across the mutagenesis population (**Table 1, Figure S3C**). The distribution and density of mutations correlated well with the genes on each chromosome, suggesting a uniform distribution of mutations (**Figure 3C**).

**Table 1.** Characterization of mutations in the KN9204 EMS-mutagenesis library.

| **Mutations characteristics** | **M<sub>2</sub>** | **M<sub>3</sub>** | **Whole population** |
| --- | --- | --- | --- |
| Number of lines | 694 | 1,396 | 2,090 |
| High-quality clean reads per line (Q30 > 0.85) | 86.44 | 94.19 | 91.62 |
| Mapping rate | 80.45% | 83.21% | 82.22% |
| Coverage of designed probe region | 99.93% | 99.93% | 99.93% |
| Average depth of coverage | 11.32 × | 11.79 × | 11.63 × |
| SNP number | 707,481 | 2,264,126 | 2,971,607 |
| Average EMS-type SNPs per line | 991.45 | 1,577.41 | 1,382.84 |
| Proportion of EMS-type SNPs | 97.26% | 97.26% | 97.26% |
| Heterozygous/homozygous ratio | 0.55 | 0.36 | 0.40 |
| Average EMS-type SNP per kilobase (population) | 3.79 | 12.35 | 19.94 |
| Mutated genes (with ≥ 1 SNP) | 102,863 | 108,725 | 108,990 |
| Mutated genes with ≥ 1 missense mutation | 81,756 | 103,140 | 106,277 |
| Mutated genes per line | 837 | 1,331 | 1,168 |
| Severely mutated gene (with ≥ 1 HIGH impact SNP) | 13,185 | 34,104 | 41,266 |
| Severely mutated genes per line | 21.32 | 33.06 | 28.90 |
| SNPs per gene model | 6.88 | 20.82 | 27.26 |
| Mutant lines per gene | 5.28 | 17.03 | 22.31 |
| Triads with ≥ 1 gene mutated | 12,975 | 12,988 | 12,988 |
| Triads with ≥ 2 genes mutated | 12,804 | 12,983 | 12,984 |
| Triads with all three genes mutated | 11,715 | 12,892 | 12,911 |
| Triads with with ≥ 1 gene severely mutated | 4,309 | 8,567 | 9,493 |
| Triads with ≥ 2 genes severely mutated | 817 | 3,851 | 5,068 |
| Triads with all three genes severely mutated | 88 | 999 | 1,589 |
| Mutant lines per triads ( with ≥ 1 gene mutated) | 16.11 | 53.05 | 69.16 |
| Mutant lines per triads (all three genes mutated) | 0.45 | 1.43 | 1.88 |

**Figure 3.**
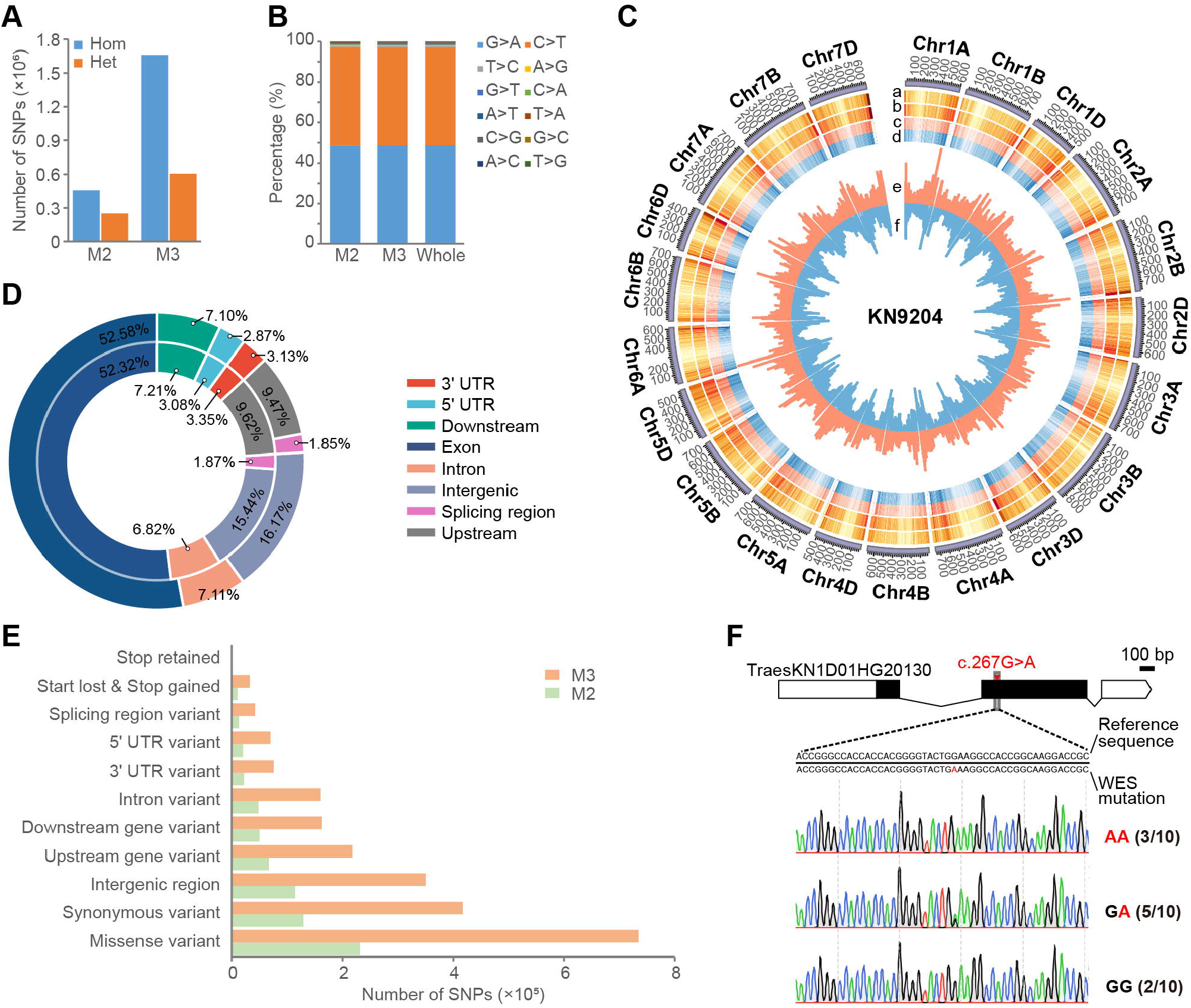
Characterization of mutations in KN9204 EMS-mutagenesis library. (A) The number of homozygous and heterozygous single nucleotide polymorphism (SNP) mutations in the M_2_ and M_3_ population. Hom, homozygous; Het, heterozygous. (B) The proportion of nucleotide mutations in M_2_, M_3_ and the whole mutagenesis population. EMS-type mutations (C>T and G>A) accounted for more than 97% SNP. (C) Circos plot of important mutation indicators in the whole EMS mutagenesis population. From the outer to the inner circle is: a. Density of probe, b. Mutation density on exon, c. The density of heterozygous mutations (light red), d. The density of homozygous mutations (light blue), e. The number of severely mutations(orange), and f. The number of SNP on the chromosome (light blue). (D) The distribution of SNPs on gene structure, with mutations of M_2_ population plot in the inner circle and M_3_ in the outer one. More than half of the SNPs were located in the exon in both populations. (E) Functional annotation of SNPs caused mutations in genes. The mutations of M_2_ population were marked in green and M_3_ in orange. (F) Validation of WES identified EMS mutations via Sanger sequencing. Numbers indicate the frequency of specific nucleotide from multiple sequencing results.

To further understand the variation patterns within structural genes, SNPs were categorized and associated with gene structure using SnpEff (Cingolani et al., 2012). The majority of variations were located in the genic region (**Figure S3D**), with approximately 52.58%, 6.82%, 6.00% and 1.85% SNPs in M_2_ population were annotated on exons, introns, UTR, and splicing region, respectively (**Figure 3D**). Functional annotation assigned 1.44%, 2.23%, and 32.50% SNPs causing stop gain/loss/retained, mis-splicing, or non-synonymous protein-coding (**Figure 3E**). To assess the credibility of mutation sites detected by WES and the transmission efficiency between generations, 22 mutations were randomly chosen for genotyping by Sanger sequencing (**Figure 3F**). In 10 homozygous mutations, 8 mutations (80.00%) were confirmed to be positive. 10 out of 12 heterozygous mutations (83.33%) were detected in a homozygous or heterozygous manner, and 2 mutations were not detected (**Table S3)**.

Thus, abundant SNP were captured in WES sequencing, with majority of them (97.26%) are reliable EMS-type mutations and retained in the progeny seeds, indicating an effective detection of high frequency mutation in KN9204 mutant library.

### Identification of gene-index mutations in KN9204 mutant library

Next, we wonder how many mutations of genome wide were actually present in KN9204 mutant library. In total, there was 2,383,763 mutations covered 108,990 genes and represented 98.79% of the annotated high confidence protein-coding genes (**Table 1**). Among them, 106,277 contained at least one missense mutation, and 41,266 genes had at least one high impact mutations (stop gained, start loss, splice donor/acceptor variant and stop retained, etc.) (**Table 1**). There are 1,168 genes mutated per line, among which 28.90 genes were severely mutated (with at least one high impact mutation), with an average of 27.26 SNP on each gene model (**Figure S3E-G, Table 1**).

We wonder whether the typical lines with visible phenotypic variations have more mutations. A similar number of SNPs (typical lines vs residual lines, 1,398.79 vs 1430.58) were observed in these lines (**Figure 4A**), but it seems that fewer genes were mutated in typical lines (1101.36 vs 1175.44) compared with residual lines (**Figure 4B**). Noteworthy, a markedly reduced number of severely mutated genes per line (28.45 vs 29.44) were observed (**Figure 4C**), suggesting that these striking mutated phenotype don’t appear to be the cumulative effect of mutated SNPs, but rather due to the mutation of pivotal genes. Importantly, 42 non-synonymous novel mutations including three stop-gained mutations (p.G247*, p.G386*, and p.G527*) were detected in the “Green Revolution” gene *Rht-D1* (Li et al., 2013) (**Figure 4D**). Four mutant lines for these three mutations exhibited marked plant height changes, with significant increased PH in homozygous lines and a separation of PH for heterozygous ones (**Figure 4E, Figure 4F**). Induced novel mutations were also detected on other genes, such as *TaTB1, TaQ, WFZP, Ms1, TaNAC019, TaCol-B5*, and *TaGW2*, etc.(Dobrovolskaya et al., 2015; Wang et al., 2017b; Dixon et al., 2018; Li et al., 2019; Liu et al., 2020a; Zhang et al., 2022; Liu et al.; Zhang et al.), among which many mutation sites have not been reported before (**Table S4**). Thus, the generated KN9204 EMS-mutagenesis library contains novel allelic variations on yield and agronomical-traits related crucial genes and holds the potential for future breeding applications.

**Figure 4.**
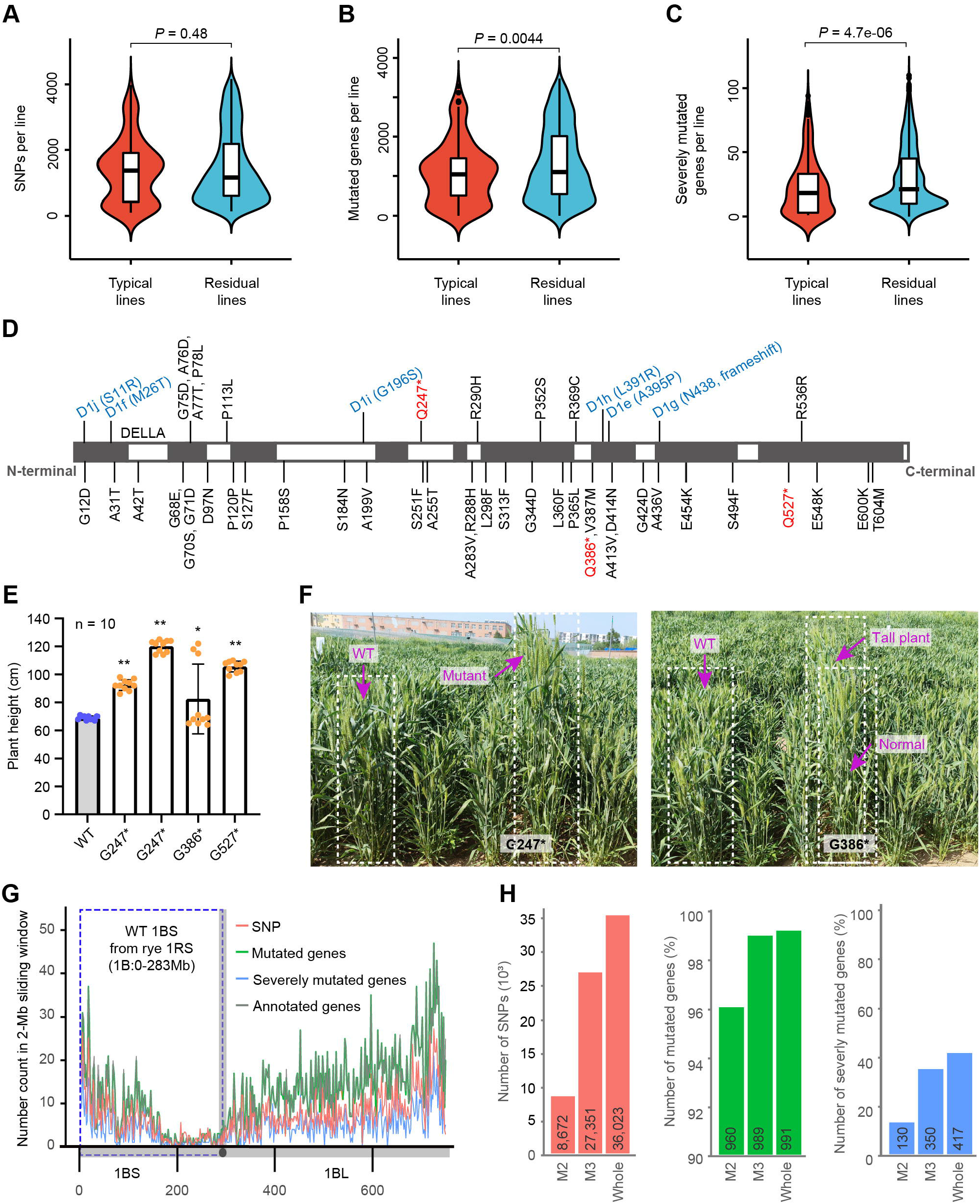
Novel allelic variations identified in KN9204 mutant library. (A-C) The comparison of SNPs per line (A), mutated genes per line (B) and severely mutated genes per line (C) of typical lines with visible phenotypic variations and the residual lines. Two-way ANOVA was used to determine significant differences. (D) Non-synonymous novel mutations detected in *Rht-D1* (*TraesKN4D01HG04200*). Schematic structure of *Rht-D1* and the positions of reported (in blue font) and novel non-synonymous mutation (in black, and stop-gained mutations marked in red). (E, F) The plant height of typical mutant lines for *Rht-D1*. The comparison of four mutant lines with wild-type KN9204 (E) and photograph of two mutant lines (F) were displayed. Note: G386* is heterozygous and the containing mutant line had plant height trait segregation. The bar plot shows the plant height of four mutant lines and wild-type KN9204, two-way ANOVA was used to determine the differences between each line with wild-type. *, *P* <0.05; **, *P* <0.01. (G). Massive mutations detected on the chromosome 1RS/1BS of KN9204. The density of SNP (in red), annotated genes (gray), mutated genes (green), and severely mutated genes (blue) were displayed on a 2-Mb sliding window (G). The blue dash box indicated the rye 1RS syntenic region (Chr1B:0-283 Mb). (H) The number of SNP, the percentage of mutated genes and severely mutated genes of EMS population in the Chr1B:0-283 Mb region. The number in the column indicates the SNP number, mutated gene number and severely mutated gene number in Chr1B:0-283 Mb region.

The common wheat is an allohexaploid species with three subgenome ((Ramríez-González et al., 2018; Shi et al., 2022). Most synteny-pair homologous genes are functionally redundant, with some homologous genes differentiate in expression pattern and function (Ramríez-González et a l., 2018). Given that, the distribution and frequency of EMS-type mutations were further evaluated using syntenic homoeolog triads as a unit. Of the 12,988 triads, mutation in at least one homologous was detected in KN9204 mutant library, with all three homologous mutated accounted for a proportion of 99.41% (**Figure S3H**). 73.09% triads were with high impact mutations in at least one homologous, and 12.23% triads were severely mutated in all three homologous (**Table 1**). For one triad, averagely 69.16 lines are accessible in the population with at least one homologous mutated, and 1.88 lines with at least one homologous severely mutated (**Table 1**). Altogether, these mutations lay a foundation for functional genomics study.

The chromosome 1BS of KN9204 was substituted by the 1RS of rye in breeding, originated from two KN9204 parents (Jimai 38 and Mianyang 75-18) (**Figure 1A**). We further explored the mutagenesis induced variation on the rye-originated 1BS in KN9204. There were fewer SNPs, mutated genes and severely mutated gene on 1BS, compared with chromosome 1BL, especially in the Chr1B:180-280 Mb that near the centromere region (**Figure 4G**). A total of 36,023 SNPs detected in the rye 1RS syntenic region (Chr1B:0-283 Mb) (Shi et al., 2022), covering 991 genes (98.41% of annotated genes), and 417 genes (41.41%) were severely mutated (**Figure 4H**). Therefore, mutants for a large proportion of genes on 1BS are still accessible in KN9204 mutagenesis library. For instance, 57 SNPs were detected in nitrate transporter *TraesKN1B01HG09350*, including a stop-gained mutation (p.Gln386*) (**Table S4**). The *ω*-secalin proteins, a main causative factor for poor quality of 1BL/1RS translocation lines, are encoded by the *Sec-1* locus on 1RS (Li et al., 2021a). We obtained several mutants of *TraesKN1B01LG01770* (**Table S4**), a secalin gene at *Sec-1* locus, which have the potential to improve the end-use quality of KN9204.

To summarize, we have built a near complete genome-wide gene-indexed mutation library, which would provide a valuable resource for genes functional analysis and may provide desirable allelic variants for breeding.

### Explore KN9204 mutant lines of NAC transcription factors for abiotic stress response

Transcription factors (TFs) are central regulators of plant development and responses to environmental stimulus (Licausi et al., 2013; Phukan et al., 2016; Kaufmann and Airoldi, 2018; Leng and Zhao, 2020), modulating broad genes expression by binding to cis-elements of the targets. By integrating PlantTFDB, HMMsearch and BLASTP, we systematically identified 11,005 TFs (**Figure S4A, Table S5**), accounting for 10.20% annotated high confidence genes, and categorize to 57 families (**Figure 5A**). WRKY, NAC and B3 are the three biggest TF families, with more than 1,000 TFs of each (**Figure 5A**). More than half of the TFs were gained along with whole genome duplicates or duplication of large genome segments (**Figure S4B**). Similar to previous reports (Evans ect al., 2022), many TFs retained in the two steps of polyploidization during the process of wheat speciation (**Figure S4C**).

**Figure 5.**
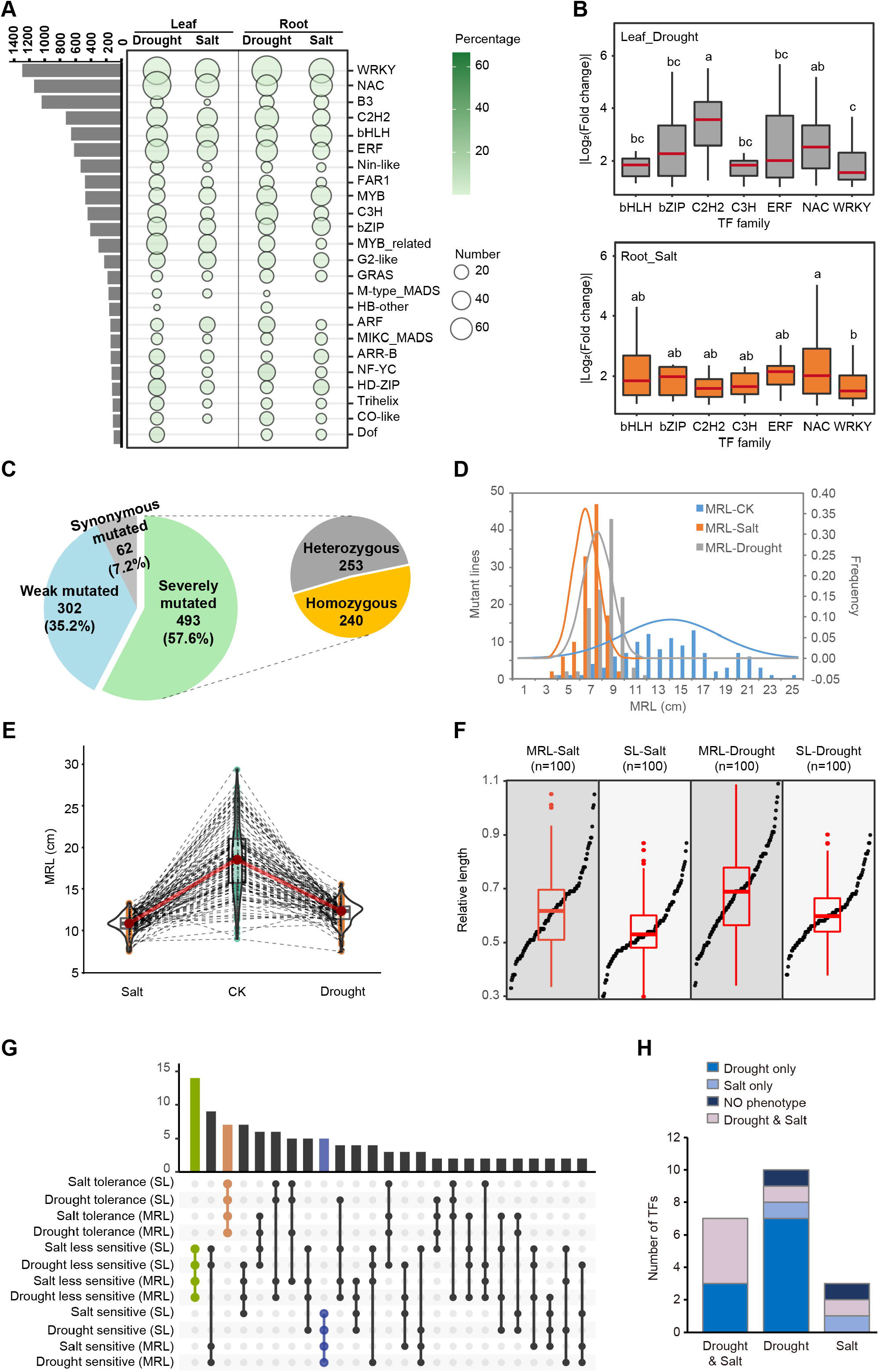
Characterization and evaluation of abiotic stress tolerance of NAC transcription factors mutant lines. (A) The number of identified TFs and the responsive of each TF family to abiotic stress. TF families with more than 100 genes were shown in the gray bars (left). The size of the circle refers the number of genes differentially expressed in root or leaf of KN9204 under drought/salt stress, with the higher percentage in darker colors. (B) Box plots show the expression level change of TF families with large number of DEGs. Upper panel, leaf under drought condition. Fisher’s Least significant difference (LSD) were used to determine the significance of expression fold change differences among TF families. Different letters mean significant difference at *P* < 0.05. (C) Summary of obtained KN9204 EMS mutant lines for NAC TFs mutations. (D) The maximum root length (MRL) distribution of 100 mutant lines under different conditions. (E) The mutant lines exhibited a distinct sensitivity to drought and/or salt stress. Each mutant line was plot in dots and their MRL under different conditions linked with a dash line. (F) The ranking of 100 mutant lines using MRL_Salt/CK_, SL_Salt/CK_, MRL_Drought/CK_ and SL_Drought/CK_. In each box, mutant lines were sort from the smallest to largest, and lines in the upper quarter, middle half and lower quarter were defined as tolerant, less-sensitive and sensitive, respectively. (G) The overview of mutant lines with different responses to drought and salt stress. (H) The number of NAC TFs response to different stress in each DEG group. The TF with at least one lines exhibited sensitive or tolerant phenotype under drought and/or salt stress were counted. The TFs differential expressed under drought, salt and both stress was assigned to the drought DEGs, the salt DEGs and drought & salt DEGs, respectively.

Abiotic stress, such as drought, salinity and heat, are threatens to wheat production and food security (Minhas et al., 2017; Shahzad et al., 2021). TFs play a critical regulatory role in response and adapt to different environmental stresses (Kaufmann and Airoldi, 2018; Leng and Zhao, 2020).

Transcriptome analysis of KN9204 seedling under drought and salt stress indicates that WRKY, NAC and bHLH TF families were abundant within the differentially expressed genes (DEGs) (**Figure 5A**). The DEGs in NAC family seems to have more dramatic changes, especially in root under salt stress (**Figure 5B, Figure S4D**). In particular, TaNAC071-A/B/D and some other NAC TFs with homologous being reported to participant in the drought, salt, heat and cold tolerance (Mao et al., 2021), were differentially expressed under stress conditions in KN9204 (**Figure S4E**), indicating potential roles of NAC TFs in response to drought and salt stress in wheat. Thus, NAC TFs containing mutant lines from KN9204 mutant library were further surveyed to explore their role in abiotic stress resistance.

A total of 1,138 NAC TFs were identified in wheat, which could be clustered into three branches based on the phylogenetic tree including wheat, *Arabidopsis thaliana* and rice (**Figure S4F**). Within KN9204 mutant library, 857 NAC TFs was identified with SNPs. Among these, 57.60% TFs with at least one high impact mutation were detected (**Figure 5C**). A total of 240 TFs get homozygous mutant lines, with a majority of TFs random mutated in few individuals (≤2) (**Figure 5C**). A total of 100 homozygous mutant lines for randomly selected 80 severely mutated NAC TFs were conducted to salt or drought treatment (**Figure S5A**). Based on pre-test in KN9204 (**Figure S5B**), 250 mM NaCl was used for salt treatment, while 8% m/v PEG-6000 was used for drought treatment. Nine indicators were measured quantitatively, among which seedling length (SL), and maximum root length (MRL) showed repeatability and proper response to salt/drought stress (**Figure S5C-E**), as reported previously (Lin et al., 2019; Ahmed et al., 2019). Both SL and MRL decreased significantly under drought and salt treatment (**Figure 5D, Figure S5F, Table S6**). However, a distinct sensitivity to drought and/or salt stress were observed among the 100 mutant lines (**Figure 5E, Figure S5G**).

We sort the MRL_Salt/CK_, SL_Salt/CK_, MRL_Drought/CK_ and SL_Drought/CK_ from the smallest to largest, and assign the upper quarter, middle half and lower quarter lines to the tolerant, less-sensitive and sensitive group, respectively (**Figure 5F**). Integrating these indexes, we assigned 100 NAC mutant lines into different groups, according to their responses to drought and salt stress. (**Figure 5G**). Among the 80 NAC TFs, 20 genes were DEGs in KN9204 under drought and/or salt conditions, correspond to 34 mutant lines. Of note, 15 TFs out of 17 DEGs (21 out of 26 mutant lines) under drought condition exhibited to be drought-sensitive or drought-tolerant, among which seven out of ten drought DEGs and all of six DEGs in both drought and salt conditions were with drought-related phenotype (**Figure 5H**). Similarly, six out of ten salt DEGs were with salt-sensitive or salt-tolerant phenotype (**Figure 5H**).

Therefore, integrating the transcriptome dynamic of KN9204 under stress treatment with KN9204 mutant library, we systematically uncovered dozens of NAC TF containing mutant lines with altered response to drought and/or salt stress.

### Uncovering the transcriptional regulation for NAC TFs in response to drought and salt stresses

Among the NAC TF mutations containing lines, detailed phenotypic observation was conducted for three lines with changed drought and/or salt sensitivity, including C332, D47, and C797 (**Figure 6A**). At CK condition, no obvious difference is observed between KN9204 and mutant lines (**Figure 6A-6C**). C332 line was sensitive to both drought and salt stress (**Figure 6A**), with significantly reduced MRL, SL, dry weight of root (DW-Root), leaf (DW-Leaf) and relative water content (RWC) compared to KN9204 (**Figure 6B, 6C**). Similar to C332 line, C797 line was sensitive to drought for both root and leaf, however its sensitivity to salt stress only in root but not for leaf morphology, such as seedling length, DW-Leaf and RWC (**Figure 6A-6C**), suggesting differential influence of C797 mutation on root or leaf tissue. On the contrary, D47 line was generally resistant to both drought and salt stress as compared to KN9204 (**Figs. 6A-6C**). Furthermore, detached leaves of D47 exhibited resistant water loss rate, implying the strong water holding capacity was detrimental to its drought or salt tolerance (**Figure 6D**). Whereas, C332 and C797 lines didn’t show difference as compared to KN9204 for water loss rate, suggesting the sensitive to drought or salt may not due to the water holding capacity change (**Figure 6D**).

**Figure 6.**
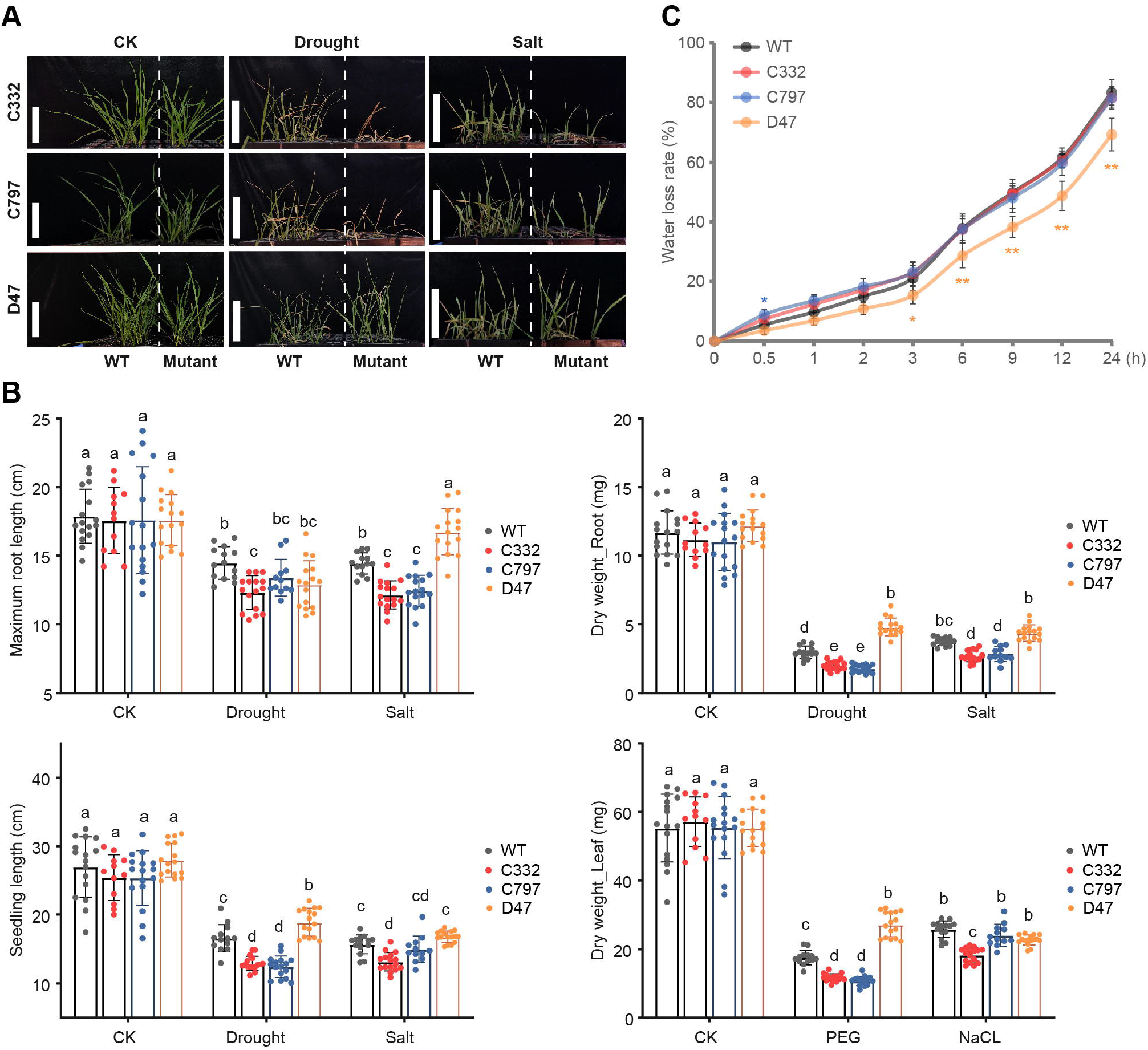
The alteration of drought and salt stress response in three NAC TF mutant lines. (A) Assessment of drought and salt tolerance of NAC TF mutant lines C332, C797, D47. Photographs were taken using seedlings two weeks after hydroponic growth under normal conditions (CK), drought treatment and salt treatment. (B) Quantitative analysis of max root length (MRL), seedling length (SL), dry weight of root (DW-Root) and dry weight of leaf (DW-leaf) in three NAC TF mutant lines under different conditions. The error bars denote ± SD, n = 15. Fisher’s Least significant difference (LSD) were used to determine the significance of trait differences among four lines (three mutant lines and wild-type KN9204). Different letters mean significant difference at *P* < 0.05. (C) The water loss rate of isolated leaves under normal condition. The fresh leaves were isolated from seedlings hydroponic growth for two weeks under normal condition. The error bars denote ± SD, n = 10. The significance of water loss rate in three mutant lines compared with wild-type KN9204 was determined by two-way ANOVA. Only those reached the significance threshold are labeled (with the color corresponding to the lines). *, *P* <0.05; **, *P* <0.01.

To understand how NAC TFs regulate abiotic stress response, we performed RNA-seq analysis for the root and leaf tissue of KN9204 and the three mutant lines (C332, C797, and D47) under control and salt or drought treatment (**Figure S6A**). Root and leaf samples are clearly separated by PCA analysis (**Figure S6A**). Within the same tissue, samples could generally be further divided by different treatments (**Figure 7A, B**), as drought treatment show more diverse than salt treatment compared to control for root (**Figure 7A**), whereas more diverse of salt treatment for leaf (**Figure 7B**). The DEG number between control and stresses further confirmed such trends (**Figure 7C**). Interestingly, D47 line is separated from other two mutant lines C332 and C797 (grouped by dashed rectangles), KN9204 is between those two groups (**Figure 7A, 7B**). This overall transcriptome pattern fits well with the resistance of D47 to salt or drought stress, while sensitive of C332 and C797 to both stresses as compared to KN9204 (**Figure 6**). Further, we compared DEG induced by drought or salt stress in KN9204 with NAC TF mutant lines, defining the altered DEGs as potential genes that regulated by NAC TFs (**Figure S6B, Table S7**). Generally, there are more shared DEG for the same tissue with different treatments than same treatment of different tissue in all three mutant lines **(Figure 7D, Figure S6C)**.

**Figure 7.**
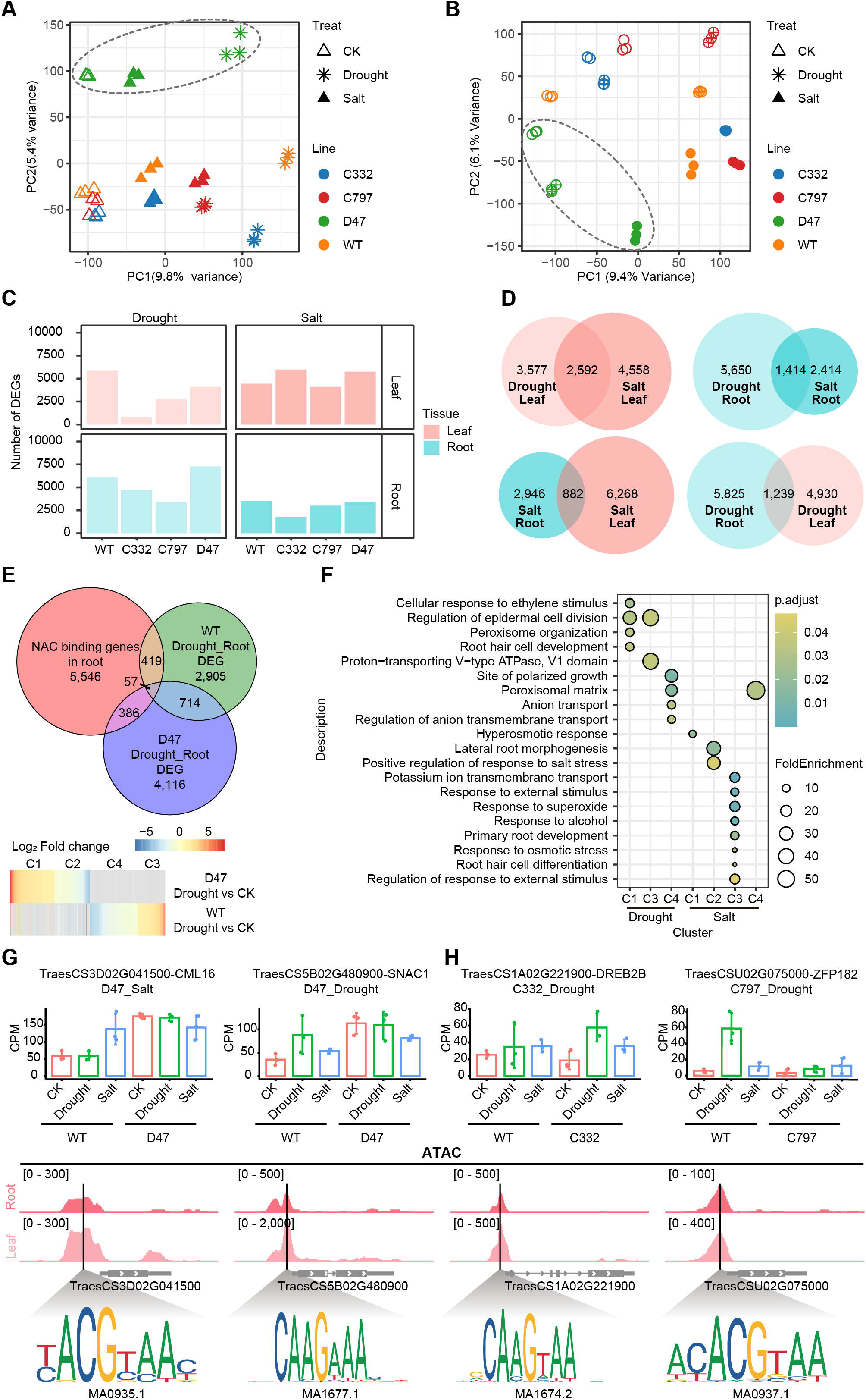
Transcriptional regulation of NAC TFs targets in response to drought and salt stresses. (A,B) Principal component analysis (PCA) of transcriptome showing the root samples (A) and leaf samples (B) under different conditions. Each dot represents one sample. Samples of different mutant lines, tissue and treatment were displayed in different shapes and colors as indicated. Dashed rectangles separate out the samples of D47 line from others. (C) Number of DEGs in root and leaf of each mutant line under different conditions. (D) The Venn diagram showing the overlapping of salt or drought stress-responsive DEGs of different tissues in mutant line C332 as compared to control. (E) Schematic of the strategy for identification of candidate targets of TraesCS5A02G242600 (D47). The KN9204 (WT) and D47 mutant line specific drought responsive genes, and genes with differential responsive level (Log_2_(FC_KN9204_/FC_mutant_)>1) summed as TraesCS5A02G242600 (D47) affected drought responsive genes. Among them, genes with NAC TF binding sites in open promoter region were considered to targets of NAC TFs during drought treatment. The heatmap shows the mRNA fold change of identified 862 drought-responsive candidate targets of TraesCS5A02G242600 in root of WT or D47 mutant line. (F) Enriched GO terms of the identified candidate targets of TraesCS5A02G242600 (D47) under drought or salt stresses. (G,H) The expression pattern of different NAC TFs candidate targets *TaSNAC1-D, OsCML16* (G)and *OsDREB-2B, OsZFP182* (H) under CK, drought or salt conditions in WT and various mutant lines (upper panel). Those candidates harbor different NAC TF binding motifs under the ATAC-seq peak in the promoter regions in root or leaf tissue, as indicated (lower panel).

Integration of chromatin accessibility data generated in KN9204 root and leaf (**Figure S1**) with the potential binding motifs of NAC TFs (**Figure S6D**) (Castro-Mondragon et al., 2022), we identified potential direct targets of NAC TFs in root and leaf (**Figure S6E, Table S8**). Majority of the potential NAC binding motifs was located in less than 1-kb distance from TSS (**Figure S6F**), and more NAC TF binding motifs were detected in root potential NAC targets (**Figure S6G**). Overlapping the NAC potential targets with NAC regulated genes by RNAseq analysis, we identified 862 drought-responsive and 774 salt-responsive candidate targets for TraesCS5A02G242600 (D47 line mutation) in root and leaf, respectively (**Figure 7E**). GO analysis showed that “cellular response to ethylene stimulus”, “root hair cell development” and antioxidation-related genes are enriched, consistent with morphological change of mutant line D47 (**Figure 7F**). In the drought-responsive TraesCS5A02G242600 candidate targets, we found wheat homologous of *SNAC1, OsLG3, OsTIFY3, OsGRAS23* and *OsXXT1*. Whereas, homologues of *OsCML16, SNAC2*, and *DSM2* were identified as the salt-responsive TraesCS5A02G242600 candidate targets (**Table S8**). Wheat homologues of *SNAC1* and *OsCML16* contain NAC binding motifs in the accessible promoter regions, showing altered induction expression in D47 as compared to KN9204 under salt or drought stress (**Figure 7G**). *SNAC1* was reported to alter the root architecture and enhance drought and salt tolerance in rice (Hu et al., 2006). Whereas for TraesCS5A02G241900 (C332) and TraesCS2D02G576400 (C797), several hundreds of candidate targets are identified for salt and drought stress (**Figure S6H**). Among them, *DREB2B* and *ZFP182* are potential targets worth to further characterize (**Figure 7H**).

Thus, taking advantage of the KN9204 mutant library and generated multi-dimensional omic data in KN9204 background, we are able to do gene functional study in an efficient way to identify NAC TFs targets that are involved in regulating drought and salt stress response/adaptation.

## Discussion

### Indexed KN9204 EMS mutant library provides important resource for gene identification and genetic manipulation in winter wheat

Bread wheat is a major crop that feeds 40% of the world’s population. Winter wheat accounts for about 70% of the total wheat acreage, with a higher grain yield, better quality, and wider adaptability than spring wheat (Entz and Fowler, 1991; Kadar et al., 2018). Gene functional study and genetic manipulation is continually undergoing for wheat, especially winter growth variety. Artificial mutagenesis does increase the genetic diversity and provide tools for gene functional study (Krysan et al., 1999; Jeong et al., 2002). Several EMS-mutagenesis populations are available in winter wheat (Chen et al., 2012; Chen et al., 2021b; Li et al., 2022). However, lack of genome wide gene-indexed information makes it hard to associate the mutant phenotype with causal DNA variation.

Here, we described a whole-exome capture sequencing characterized EMS mutagenesis population in winter wheat variety KN9204 (**Figure 1, Figure 3**). Recently, we have assembled a high-standard reference genome of KN9204 (Shi et al., 2022), and designed the whole exome capture probes accordingly. By contrast, the current existing EMS mutant libraries don’t have high standard reference genome, a pseudogenome was assembled in spring wheat variety Cadenza (Krasileva et al., 2017), while others are not sequenced, such as Lunxuan987 and Jinmai47 (Chen et al., 2021b; Li et al., 2022). The lack of genome sequence brings inconvenience to the utilization of mutations, such as the retrieval and selection of desired mutants, the primer design and authentication of variations, etc.

Importantly, KN9204 mutant pool also has advantages in the mutation frequency and pool size, two key parameters that determine the available variability of the mutagenesis population (Henry et al., 2014). A high mutation density of at least one mutation per 50.2 bp was obtained in the KN9204 population, slightly less than that reported in tetraploid Kronos (∼28.7 bp), and hexaploid Cadenza (∼25.3 bp) (Krasileva et al., 2017), but much higher than that of Longfumai 17 (∼244.2 kb) (Li et al., 2017), NN-Gandum-1 (∼47.8 kb) (Hussain et al., 2018), Jinmai47 (∼47 kb) (Chen et al., 2012), Zhongyuan 9 (∼0.6 kb) (Li et al., 2022), and *T. monococcum* accession TA4342-96 (∼92 kb) (Rawat et al., 2012). With 2090 lines, KN9204 mutant library provides variations covering the nearly entire genome coding genes (98.79%) and produce severe mutations for 41,266 high confidence genes (**Figure 3, Table 1**). The 1RS chromosome segment from rye has been widely used in the breeding programs for agronomic traits improvement (Feuillet et al., 2008; Moskal et al., 2021). KN9204 is the only published wheat variety with genome carrying 1BL/1RS wheat-rye translocated chromosome to date (Shi et al., 2022). At the 1RS chromosome region, we detected mutation for 98.41% annotated genes and novel variation on *ω*-secalin protein genes (**Figure 4**), which could be useful for understanding the genetic basis for elite trait formation and breeding improvement.

We did in-field survey for visible developmental alterations and observed diverse morphological variations in KN9204 mutant population, covering important agronomic traits (heading date, plant height and leaf morphologies), yield traits (SNPS, TGW and GNPS), quality (grain hardness, suitability) (**Figure 2**). More importantly, via WES we identified novel allelic variations on agronomic- and yield-related genes, such as *Rht-D1, WFZP, GW2*, etc. For the case of “Green Revolution” gene *Rht-D1*, we detected three novel alleles with truncation in multiple lines, of which four lines showed altered plant height **(Figure 4, Table S4**). Thus, the genetic variation generated from KN9204 mutant library holds the potential for breeding process.

Therefore, a large-scale mutant library with abundant genetic variation was developed in winter wheat KN9204, an elite variety with high quality reference genome carrying 1BL/1RS wheat-rye translocated chromosome, is expected to play important role in functional genomics and breeding.

### All-In-One Toolbox of KN9204 for gene functional study in wheat

Recent years, numerous genomic study and high-throughput multi-omic sequencing data enabled wheat research into a “big data era” (Xiao et al., 2022). Integration of different layers’ omic data could accelerate the understanding of wheat productivity, quality and environmental adaptation. Because of the elite agronomic trait and high quality reference genome of KN9204 (Jia et al., 2006; Cui et al., 2016; Shi et al., 2022), we have generated transcriptome and epigenomic dataset including chromatin accessibility, core histone modification and variants from multiple tissues, developmental stages and environmental treatments (**Figure S1**) (Li et al., 2018; Zhao et al., 2022; Lin et al., 2022; Shi et al., 2022; Zhang et al., 2022). These data combined with the KN9204 gene-indexed mutant population established here would propel the wheat functional genomics at different research levels.

NAC TFs are known to regulate abiotic stress especially for drought resistance in wheat and other species (Jeong et al., 2010; Mao et al., 2015; Mao et al., 2020; Chen et al., 2020; Mao et al., 2021). In this study, through integrating the transcriptome dynamic of KN9204 under stress treatment with KN9204 mutant library, we systematically surveyed 100 mutant lines of 80 NAC TFs and uncovered 13 NAC TF containing mutant lines with altered response to drought and/or salt stress (**Figure 5**). Such medium-scale screening and verification strategy is also applied by us for wheat spike development(Lin et al., 2022). We screened 85 potential TFs, which is identified by integration of transcriptional regulation network with GWAS, and found 44 TFs containing KN9204 mutant lines have altered spike architecture (Lin et al., 2022). With QTL data generated by KN9204-Jing 411 RIL population for various agronomic traits (Fan et al., 2018; Fan et al., 2019; Liu et al., 2020b; Yu et al., 2022; Zhao et al., 2022), we would expect that verification and genetic effect evaluation of candidate genes located in the interval of QTL from KN9204 can be easily performed via KN9204 mutants. Thus, the KN9204 EMS population assists the mining, identification and verification of genes underlying important traits.

Moreover, combining multi-omic data generated in KN9204 **(Figure 1**) with indexed KN9204 mutant lines can accelerate the interpretation of molecular regulatory mechanisms for specific genes, especially TFs. Here we demonstrated a snapshot of transcriptional regulatory analysis of three NAC TFs participating in drought and/or salt response/adaptation (**Figure 6, Figure 7**), by integrating transcriptome, chromatin accessibility and TF binding sites resources previously generated in KN9204 (Shi et al., 2022; Zhang et al., unpublished). These multidimensional omics data resources generated in KN9204 conducive to the study on effects of DNA variation on gene expression, transcriptional regulatory network, and potential interaction proteins, etc.

To summarize, we provide a “All-In-One” toolkit in winter wheat variety KN9204 background for the wheat research community, and hope it could trigger the “fast forward key” for wheat functional genomics and prosperity of wheat mutant breeding.

## Materials and Methods

### Plant material and EMS mutagenesis

Winter wheat variety KN9204 was used to generate the mutant populations. To ensure the purity and uniformity of seeds, grains of ten plants derived from one spike were used for mutagenesis. In brief, approximately 5,000 seeds (M_0_) were pre-soaked in ddH_2_O for 6 hours, then soaked and gently shaken for 12 hours in 1.0% (v/v) EMS (liquid, 99% purity; Coolaber, China) solution at the ratio 200 grains/100 mL at room temperature (25℃). The treatment was terminated by washing the seeds for 10 minutes in 200 mM sodium thiosulfate solution and then thoroughly washed with running water for 1 hour. The EMS-treated seeds were sown in the field, 2,090 M_1_ individuals survived to harvested M_2_ seeds. Among these, 1,396 M_2_ plants successively advanced to the M_3_ generation using a single spike in greenhouse. Sixteen progenies of the 694 M_2_ and 1,396 M_3_ individuals were planted in a row in the field for phenotypic investigation and DNA extraction, all individuals in each row were harvested and bulked for later distribution. For each row, the young leaves of eight to ten seedlings were sampled and pooled for genomic DNA extraction using the Plant Genomic DNA Extraction Kit (DP205, TianGen, China) following the instructions provided by the supplier.

### Growth phenotyping in filed

Sixteen progenies of the M_2_ and M_3_ individuals were planted in the field for phenotypic investigation. Each mutant line was space planted in a 1.5-m single-row plot with 10 cm between plants and 25 cm between rows, and managed following the local practices. Two independent surveys were conducted during the 2021–2022 cropping seasons in the fields at Beijing (39°55’ N, 116°23’ E) and Shijiazhuang (38°04’ N, 114°28’ E). Anomalous developmental defects observed in both fields and each with at least three individuals were considered credible phenotypic variations. Thirteen quantitative indicators were investigated in both fields for a randomly chosen 200 mutant lines, and the mean value of each trait were used for analysis. Flowering time was calculated as the days from sowing to flowering of half of spikes in a row. Flag leaf length and width were investigated 10 days after flowering, using main shoots of five plants in the center of each row, by a leaf area measuring instrument (Yaxin-1242, Beijing Yaxin Science Instrument Technology Co., Ltd., Beijing, China). Plant height, spike number per plant and tiller angler were measured 15 days after flowering, with ten uniform plants in the center of each row. Plant height were measured from soil to the tip of the uppermost spike (excluding awns). Spikes with at least three grains were counted to measure the spike number per plant. Tiller angle were measured with an angle measuring instrument (TPZW-J-1, TOP Inc., China). Spike length and spikelet number per spike were measured with ten main tiller spikes. Spike length was measured from the base of the spike to the tip (excluding the awns), and the total spikelet number per spike were counted. These spikes were harvested and used to measure the grain number per spike, 1000-grain weight, grain length, grain width and grain roundness, by a Crop Grain Appearance Quality Scanning Machine (SC-G, Wanshen Technology Company, China).

### Construction of exome sequencing libraries

Genomic DNA was quantified using a Nano-drop 2000 spectrophotometer (Thermo Fisher Scientific, Waltham, MA). Approximately 2 ug of genomic DNA was fragmented into 200-300 bp using the Bioruptor UCD-200 sonicator (Diagenode, Denville, NJ). Libraries were constructed using the DNA library prep kit for BGI platform (BGI, Shenzheng, China). Fragmented DNA was end repaired with an end-repair enzyme and a deoxyadenosine. The adapter-ligated libraries were selected for an average insert size of ∼350 bp using magical beads according to the manufacturer’s instructions (Twist, CA, USA). The pre-capture libraries amplification was performed with 4∼5 PCR cycles. Equal amounts of products from eight libraries were pooled to obtain a total of 4 ug of DNA for the hybridization.

Hybridization of sample libraries was performed using the customer exome panel (Tcuni, Chengdu, China), the exome panel specifically designed for the CDS of KN9204 genome, including 1,232,636 probes covering 127 Mb genome (98.11% CDS of HC gene). The hybrids were captured using the M1 (Thermofisher, MA, USA) capture beads kit, washed, and amplified by ligation mediated PCR. The quality of the captured libraries was assessed using the Agilent 4200 Tape Station (Agilent, CA, USA). Final captured libraries were sequencing with BGI T7 platform.

### Reads mapping and SNP calling

After quality control of raw reads by fastp (v0.19.5), bwa (v0.7.17-r1188) was used to align reads to the KN9204 reference genome shared by Shi et al. prior to publication (includes 110,326 genes) with default settings. Reads with MQ<30 were removed. BAM files were sorted and PCR-duplicates were removed using samtools (v1.9). GATK best practice workflow was utilized for SNP calling. A GVCF file was generated for each sample through the ‘HaplotypeCaller’. Then, ‘GenotypeGVCFs’ was used to transform the GVCF file into a raw VCF file for each sample. Finally, bcftools (v1.9) was used to filter each VCF file by ‘%QUAL<100 || INFO/DP<5’. In total, we obtained 34,156,989 SNP from all the sequenced samples.

### Identification of EMS mutations

In order to reduce the false-positive rate of EMS mutations among the SNP datasets, a strict pipeline of filtering was applied, including three steps, 1) 614,725 SNPs were discovered among three wild-type plants of KN9204. Those SNPs were caused by errors of sequencing or computational processing; therefore, they were removed from our dataset. 2) EMS mutations among a large genome were rare events. It is unlikely a genomic site being mutated in more than one sample. So, any SNP detected in more than one samples were considered as errors. 13,862,902 SNPs were remained in our dataset. 3) We removed heterozygous SNPs supported by less 10 and homozygous SNP supported by less than 6 reads. As a result, 2,971,607 SNPs were defined as our final EMS mutations and used in our analysis. Since users may want to retrieve all possible mutations, a larger SNP dataset were used in our website, where one mutation was allowed to present in no more than 4 samples.

### Validation of EMS mutations

A total of 22 mutations were randomly chosen for validation through direct Sanger sequencing. The main objectives were to confirm the presence of the mutation, and assess the mutation transmission efficiency in progeny seeds. For each mutation, At least four plants were used for validation, the gene specific PCR primers used for cloning genomic regions flanking the target mutation were shown in supplemental **Table S3**.

### Access to the mutation and seed stocks

The EMS-type mutations detected in the KN9204 populations at different stringency levels are accessible in public databases (http://183.223.252.63:9204/) and can be visualized using a JBrowse graphic interface, the mutations were colored based on severity of mutation effects (red = stop gained/frameshift, violet = missense, green = synonymous, and blue = non-coding regions). Additional mutation information is available when right-clicking on a particular mutation in the browser. “Blast”, “Get sequence” and “PrimerServer” tools base on KN9204 genome and genes were integrated to allow users easily get gene ID and primer of KN9024. Once the desired mutations are identified, users can make an order on “Query” page and the background system will automatically send the order to both user and manager.

### Annotation and phylogenetic tree generation of NAC TFs

We annotated the TFs by taking the union of orthologs-based results, HMMsearch approach and BLAST-based approach. HMMsearch were carried out using the pfam conserve domain of TF families (provided by PlantTFDB) in wheat genome to identify wheat transcription factors. Protein sequences of TFs annotated in rice and Arabidopsis were performed to homologous alignment with wheat genes, using BLASTP method with cutoff of e-value <= 1e^-5^. We aligned the NAC protein sequences of wheat, rice and Arabidopsis using ClustalW, and create the maximum likelihood phylogenetic trees using the auto setting to detect the best protein model, 100 maximum likelihood searchers, and 100 rapid bootstraps.

### Stress treatments

Seeds of KN9204 and the mutant lines were surface-sterilized in 1% sodium hypochlorite for 15 min, then washed in distilled water several times, and finally laid on vermiculite for 5 days. Uniformly seedlings were wrapped with sponge and fixed on a seedling-raising plate through a 1-cm hole in diameter. Prior to stress treatments, the seedlings were grown hydroponically in half-strength Hoagland solution in the green house for 24 h, then the seedlings were subjected to drought stress (8 % PEG-6000, m/v) and salt stress (250 mM NaCl) for 14 days, in a growth chamber with 22 °C/18 °C (day/night), 16 h/8 h (light/dark) and 50% humidity. All experiments were performed in parallel and seedlings in normal growth condition were taken as control. Leaves and roots were collected separately at 6 days after stress treatment and frozen immediately in the liquid nitrogen, and stored at −80 °C.

### Transcriptome analyses

Total RNA was isolated from the leaf and root under normal and stress conditions (three biological replicates were conducted for each treatment), using an RNAprep Pure Plant Kit (TIANGEN, Beijing, China). Nano-Drop 1000 spectrophotometer and Agilent Bio-analyzer 2100 system (Agilent Technologies Co. Ltd., Beijing, China) was used to determine the quantity and quality of the RNA. Equal amounts of total RNA of 20 samples were pooled for library construction and transcriptome sequencing using a modified strand-specific 3’-end RNA-Seq protocol. Raw reads were filtered by fastp v0.20.1 for adapters removing, low-quality bases trimming, and reads filtering (Chen et al., 2018). Furthermore, FastQC v0.11.8 (http://www.bioinformatics.babraham.ac.uk/projects/fastqc/) was performed to ensure the high quality of reads. Clean Reads were aligned using hisat2 v2.1.0 with default parameters (Kim et al., 2019). The number of paired reads that mapping to each gene was counted using feature Counts v2.0.1 with default parameters (Liao et al., 2014). The counts files were then used as inputs for DEGs (differentially expressed genes) analysis by DESeq2 v1.26.0 with a threshold “absolute value of Log2 Fold Change ≥ 1 and FDR≤ 0.05” (Love et al., 2014). CPM values generated from the counts matrix were used to characterize gene expression and used for PCA analysis. For subsequent clustering and visualization, we obtained mean counts by merging three biological replicates. Functional enrichment was performed using an R package cluster Profiler v3.18.1 (Yu et al., 2012), and GO annotation files were generated from IWGSC Annotation v1.1.

### Statistical analysis used in this study

R (https://cran.r-project.org/; version 4.0.2) was used to compute statistics and generate plots if not specified. The two-way ANOVA was used to perform the two groups’ comparison of data (**Figure 4A-C, Figure 4E, Figure 6C**). For three or more independent groups comparison of data, Fisher’s Least Significant Difference (LSD) was used (**Figure 5B, Figure 6B, Figure S4D**). We used Wilcoxon’s two-sample test to detect the differences of NAC TF binding sites number between leaf and root (**Figure 7G**).

## Data availability

The raw sequence data of RNA-seq and ATAC-seq reported in this study have been deposited in the Genome Sequence Archive (Chen et al., 2021a) under accession number CRA009130, in National Genomics Data Center (CNCB-NGDC Members and Partners, 2022), China National Center for Bioinformation / Beijing Institute of Genomics, Chinese Academy of Sciences that are publicly accessible at https://ngdc.cncb.ac.cn/gsa. The published RNA-seq and epigenomic data of KN9204 used for analysis is download from CNCB-NGDC under accession number CRA008877 (spike related data, Lin et al. 2022) and PRJCA004416 (leaf and root related data, Shi et al. 2022).

## Author contributions

J.X. and X.-G. L. designed and supervised the research; J.X., D.-Z.W. wrote the manuscript with help of H.-J.W. F.H. Y.-H. H. Y.-L. J. X.-G.L. J.-M. L. Y.-M. Y. Y.-Q. Z. Y.G. C.-J. Z. W.-Q. T. L.W. X.-L.G. S.-Z.Z. Y.-J.Z. X.-L.L.; J.X. Y.-M. Y. D.-Z.W. H.-J. W. F.H. Y.-X. X. prepared all the figures; Y.-P.L. F.C. X.-G.L. J.-M.L. generated the EMS mutant library; Y.-P. L. D.-Z.W. did mutant library growth in fields and phenotyping; Y.-X.Z. Z.-X.C. L.-X.G. did genome DNA extraction for WES sequencing and Sanger sequencing for verification; H.-J. W. and F.H. did WES sequencing analysis; Y.-X. X. did the transcriptome and epigenome analysis. D.-Z. W. did all the rest of experiments.

## Acknowledgements

This research is supported by the Strategic Priority Research Program of the Chinses Academy of Sciences (XDA24010204) to J.X., Hebei Natural Science Foundation (C2021205013) and ‘Full-time introduction of high-end talent research project’ (2020HBQZYC004) to X.-G. L., National Natural Science Foundation of China (U22A6009) to J.-M. L., the Research Program for Network Security and Information of the Chinese Academy of Sciences (CAS-WX2021SF-0109) to F.H. and J.X., and the National Key Research and Developmental Program of China (2021YFD1201500) to J.X.

## Figure legends

**Figure S1. Multi-omic data generated in KN9204**

(A) Various of histone modification, chromatin accessibility and transcriptome datasets of typical tissues across different developmental stages and conditions. The wheat growth stage of root and leaf was recorded using Zadok’s scale. SAM, shoot apical meristem; EL, elongation stage; SR, single ridge; DR, double ridge; SMI, spikelet meristem initiation; GPD, glume primordium differentiation; FMI, floral meristem initiation; FOP, floral organ primordium differentiation. CK, normal growth condition; LN, low nitrogen. ※, transcriptome data from Shi et al. (2022) under accession number PRJCA004416; *, The RNA-seq and epigenomic data of KN9204 spike are available from Lin et al. (2022) under accession number CRA008877; ^, unpublished data;

(B) IGV browser showing gene expression, chromatin accessibility and histone modifications of root (Zadoks12 stage), flag leaf (Zadoks55 stage) and young spike (double ridge stage) at *TaNAC071-A*.

**Figure S2. Distribution of quantitative traits in the KN9204 EMS-mutagenesis population**.

(A-I) Various quantitative indicators were investigated in a randomly chosen 200 mutant lines, as indicated. The red dash lines indicate trait value of wild-type KN9204. The frequency distribution of each trait was shown in histogram with a fitting curve (in gray). The indicators fit an approximate normal distribution were in light green (A-G), the indicator fit positively skewed distribution was in light blue (H) and the negatively skewed distribution ones in brown (J, I).

**Figure S3. Quality analysis for the exome captured sequencing of KN9204 mutant library**

(A) The proportion of high-quality clean reads reach a specific sequencing depth.

(B) The proportion changes of EMS-type SNPs in heterozygous (in blue) and homozygous (in green) mutations along with the sequencing depth. The dash lines indicate the minimum reads coverage used to identify the high credibility variations (HetMC10 and HomMC6).

(C) The SNP density in M_2_ and M_3_ populations.

(D)The distribution of SNPs on the gene region

(E, F) The distribution of mutated gene number (E) and severely mutated gene number (F) per line in M_2_ and M_3_ populations.

(G) Number of mutations for each gene model.

(H) The number of accessible mutant lines per triad with SNP (brown) and high impact SNP (blue) for at least one homologous gene.

**Figure S4. The expansion and expression pattern of NAC TFs in wheat**.

(A) Venn diagram of TFs number identified through three methods.

(B) Number and percentage of genes gained when *Triticum*. and common wheat was differentiated.

(C) The number and proportion of different gene duplication types for each TF family. Only TF families with more than 100 genes were displayed.

(D) Box plots show the expression level change of TF families with large number of DEGs. Upper panel, leaf under salt condition. Fisher’s Least significant difference (LSD) were used to determine the significance of expression fold change differences among TF families. Different letters mean significant difference at *P* < 0.05.

(E) Heatmap of NAC TFs differentially expressed under stress conditions in KN9204.

(F) The NAC TFs was clustered into three branches based on the phylogenetic tree. Clusters were marked with different colors, and protein domains shown in the phylogenetic tree different color rectangles. Bars = 100 bp.

**Figure S5. Phenotypic analysis of NAC TFs mutant lines**.

(A) A photograph showing the equipment and hydroponics growth environment of wheat seedlings for PEG and salt treatment.

(B) The plant growth curve of KN9204 seedling under different treatments, indicated by seedling length.

(C) A radar graph representing the percentage of each trait in KN9204 under different conditions. RSA, root surface area; Forks, number of root forks; Tips, number of root tips; TRL, total root length; RLPV, root length per volume; SL, seedling length; MRL, max root length; RV, root volume; RAD, root average root diameter.

(D) A ridge chart displaying the distribution of seedling length (SL) and max root length (MRL) in NAC mutant lines under different conditions.

(E) Correlation heatmaps of SL (leaf) and MRL (right) in NAC mutant lines under different condition and repeat batch. The numbers in the squares refer to pair-wise Pearson’s correlation coefficients (*R* value).

(F) The SL distribution of mutant lines under different conditions.

(G) Comparison of SL for each mutant line under different conditions.

**Figure S6. Transcriptome analysis of various NAC TFs mutant line under different conditions and identification of NAC TFs candidate targets**

(A) Principal component analysis (PCA) of all samples of NAC mutant lines C332, C797, D47 and wild-type KN9204 under CK, drought or salt conditions.

(B) Schematic of the strategy for identification of D47 affected drought responsive genes. KN9204 (WT) and D47 mutant line specific drought responsive genes, and genes with differential responsive level (Log_2_(FC_KN9204_/FC_mutant_)>1), as indicated by arrow, are summed as TraesCS5A02G242600 (D47) affected drought responsive genes

(C) Four-way Venn diagram of DEGs in root or leaf of mutant lines C332, C797 and D47 under salt or drought conditions to show the overlapping pattern.

(D) NAC TFs binding motifs were clustered into four clades based on motif similarity. A representative motif in each clade was shown below.

(E) Venn diagram showing the overlap and unique candidate targets of NAC TFs in root and leaf.

(F, G) The number (F) and distribution pattern (G) of NAC TF binding motifs in the promoter regions of NAC TFs candidate targets in leaf and root.

(H) Summary of candidate target numbers of TraesCS5A02G242600 (D47), TraesCS5A02G241900 (C332) and TraesCS2D02G576400 (C797) under salt and drought stress in leaf and root.

## Supporting information

Table S1 Anomalous developmental mutants of each category in KN9204 EMS mutation population. Table S2 Quantitative traits performance of KN9204 and range in EMS mutation population.

Table S3. Validation of mutations identified in WES.

Table S4 EMS-type SNPs identified in some important genes.

Table S5. Genome-wide identification of transcription factor families through different methods.

Table S6. Summary of nine quantitative indicators of the 100 NAC TF mutant lines under different conditions.

Table S7. DEGs list of mutant lines

Table S8. Genes with NAC binding motif in leaf and root

