## Supplemental Table 1 for "Boosting wheat functional genomics via indexed EMS mutant library of KN9204"

**Table S1 Anomalous developmental mutants of each category in KN9204 EMS mutation population.**

| **Category** | **Phenotype description** | **Number of**  **mutants observed** | **Frequency** |
| --- | --- | --- | --- |
| Plant architecture | Dwarf (PH < 50 cm) | 6 | 0.29% |
|  | Tall plant (PH > 100 cm) | 17 | 0.81% |
|  | Plant height separation (ΔPH > 20 cm) | 71 | 3.40% |
|  | Low-tillering (< 5) | 6 | 0.29% |
|  | Procumbent | 3 | 0.14% |
|  | Loose plant | 4 | 0.19% |
|  | Compact plant | 9 | 0.43% |
|  | Tufted | 7 | 0.33% |
|  | **Summary** | **123** | **5.88%** |
| Spike and affiliated organs | Non-terminated floret | 2 | 0.10% |
|  | Glume hair | 7 | 0.33% |
|  | Paired spikelet | 93 | 4.45% |
|  | Short awn | 10 | 0.48% |
|  | Black glume and awn | 2 | 0.10% |
|  | **Summary** | **114** | **5.45%** |
| Leaf shape and color | Chlorisis | 7 | 0.33% |
|  | Wax leaf | 23 | 1.10% |
|  | Striped leaf | 5 | 0.24% |
|  | Twisted leaf | 3 | 0.14% |
|  | Lesion mimic | 3 | 0.14% |
|  | **Summary** | **41** | **1.96%** |
| Other | Purple seed coat | 4 | 0.19% |
|  | Early or late heading (> 5 days) | 13 | 0.62% |
|  | Strong vernalization demand | 3 | 0.14% |
|  | Sterile | 2 | 0.10% |
|  | Premature necrosis | 2 | 0.10% |
|  | Premature senescence | 45 | 2.15% |
|  | Susceptible to leaf rust | 2 | 0.10% |
|  | Susceptible to powdery mildew | 3 | 0.14% |
|  | **Summary** | **74** | **3.54%** |
| **Total** |  | **286** | **13.68%** |
