## Supplemental Table 2 for "Boosting wheat functional genomics via indexed EMS mutant library of KN9204"

**Table S2 Quantitative traits performance of KN9204 and range in EMS mutation population.**

| **Phenotype description** | **Character range** | |
| --- | --- | --- |
|  | **KN9204** | **EMS population** |
| Plant height (PH, cm) | 68.90 ± 1.45 | 42.75 - 128.40 |
| Flowering time (FT, days) | 193 ± 1.03 | 184 - 198 |
| Spike length (SL, cm) | 10.25 ± 0.47 | 6.26 - 14.38 |
| Spikelet number per spike (SPS) | 25.54 ± 1.66 | 17.25-34.62 |
| Spike number per plant (SN) | 18.58 ± 3.29 | 3.60 - 27.56 |
| Tiller angler (TA, °) | 13.26 ± 0.85 | 5.22 - 35.67 |
| Flag leaf length (FLL, cm) | 21.40 ± 2.22 | 11.70 - 28.90 |
| Flag leaf width (FLW, cm) | 1.92 ± 0.12 | 1.08 - 2.78 |
| Grain number per spike | 57.84 ± 3.50 | 27.42 - 95.07 |
| TGW (g) | 46.49 ± 0.90 | 24.04 - 62.72 |
| Grain Length (mm) | 6.03 ± 0.02 | 5.36 - 7.19 |
| Grain width (mm) | 3.55 ± 0.03 | 2.64 - 4.01 |
| Grain roundness | 0.59 ± 0.01 | 0.44 – 0.63 |
